# Dysfunction of a SET3-like complex underlies a family of related neurological disorders

**DOI:** 10.1101/2025.05.30.657039

**Authors:** Katie M. Paton, Beatrice Alexander-Howden, Jenna I. Hare, Jacky Guy, Kashyap Chhatbar, Maria Yudina, Robyn Walls, Tricia Mathieson, Christos Spanos, Adrian P. Bird, Matthew J. Lyst

## Abstract

TBLR1 is a subunit of the NCoR corepressor complex that is mutated in a range of neurodevelopmental disorders. Here, we report that TBLR1 functions as a molecular scaffold that physically connects ANKRD11 and SETD5 – two of the most frequently mutated proteins in neurodevelopmental disorders – and links them to the rest of the NCoR complex. The resulting assembly resembles the yeast SET3 complex (SET3C) – a transcriptional regulator. Pathogenic missense mutations in TBLR1, ANKRD11 and SETD5 disrupt this assembly, and an engineered mutation that specifically abolishes SETD5 incorporation into SET3C causes severe developmental impairments in mice. Disruptions of mammalian SET3C components cause highly correlated changes in gene expression – including upregulation of already highly transcribed genes. Together, our results reveal that failure of transcriptional regulation by SET3C is a convergent molecular basis for a family of neurodevelopmental disorders.

## Introduction

Neurodevelopmental disorders often arise from *de novo* loss-of-function mutations in single genes. Mutations in any individual gene contribute only a small fraction of total cases, but the cumulative disease burden of lesions in hundreds of different genes is large^1^. Most implicated protein products fall into broad functional categories, including synapse proteins, translational regulators and chromatin proteins^2,3^. In nearly all cases, however, the chain of events linking the mutation to its phenotypic consequences is unclear. Specifically, the extent to which mutated proteins within these categories might function together in shared molecular pathways is not known. This study focusses on the nuclear corepressor (NCoR) complex and, in particular, the subunit TBLR1, encoded by the *TBL1XR1* gene. Mutations in this gene are associated with intellectual disability and developmental delay, but do not cause a single well-defined syndrome. Instead, different mutations in TBLR1 lead to distinct clinical outcomes. For example, complete deletion of TBLR1 leads to relatively mild intellectual disability^4,5^, mutations affecting the N-terminal LisH domain cause West syndrome with autistic features^6–8^, and mutations affecting the C-terminal WD repeat (WDR) domain often cause severely delayed language development, intellectual disability and gastrointestinal problems^9^. A particular mutation in the WDR domain, Y446C, causes Pierpont syndrome, a distinctive disorder characterized by a combination of intellectual disability, hearing loss, facial dysmorphism and abnormal fat distribution in the limbs^10^. TBL1X, a closely related paralog of TBLR1, is also a subunit of NCoR. Mutations in the *TBL1X* gene appear not to be associated with neurodevelopmental disorders, but instead are linked to hypothyroidism and hearing loss^10,11^.

The first clinically relevant TBLR1-interacting protein to be identified was the methyl-CpG binding protein MeCP2^12–14^, mutations in which cause the severe neurological disorder Rett syndrome (RTT)^15^. RTT is characterised by repetitive hand stereotypies, gait abnormalities, loss of speech and intellectual disability^16–19^. MeCP2 recruits the NCoR complex via a well-defined interaction with the WDR domains of its TBL1X and TBLR1 subunits^20,21^. MeCP2 itself binds broadly across the genome^22^ and evidence suggests that it brings the NCoR complex to chromatin, leading to transcriptional inhibition, in particular of highly methylated genes^23–29^. Supporting the idea that NCoR recruitment to DNA is a critical function of MeCP2^30^, separate clusters of RTT-causing missense mutations disrupt the DNA-binding methyl-CpG binding domain (MBD) or the NCoR interaction domain (NID) of MeCP2^20^. In fact, all characterised RTT-causing mutations in MeCP2 affect the ability of MeCP2 to recruit the TBLR1-containing NCoR complex to chromatin^31^, and radically truncated versions of MeCP2, preserving the MBD and the NID, rescue most of the consequences of MeCP2 deficiency in mice^32^. Given that MeCP2 function relies on its interaction with TBLR1, patient mutations in TBLR1 might be expected to disrupt MeCP2 binding and result in RTT-like symptoms. However, whilst one individual with a clinical diagnosis of RTT has a TBLR1 mutation located at the MeCP2 binding interface^21,33^, the disparate symptoms of most patients overlap only partially with RTT. For this reason, we hypothesized that many pathogenic TBLR1 mutations affect interactions and functions that are MeCP2-independent.

Here, we have investigated the molecular pathology associated with TBLR1 dysfunction. We examined the molecular consequences of pathogenic TBLR1 missense mutations, and found that, whilst many disrupt binding to MeCP2, interactions with SETD5 and/or ANKRD11 were also strongly affected. SETD5 and ANKRD11 have been reported to interact with the NCoR complex^34–37^, and, like MeCP2, are themselves among the ten most frequently mutated genes in neurodevelopmental disorders^38^. Mutations in ANKRD11 cause KBG syndrome (KBGS), which is characterized by intellectual disability, facial dysmorphism, macrodontia, and skeletal abnormalities^39^. Interestingly, the spectra of symptoms for SETD5 syndrome – intellectual disability and dysmorphic facial features – often overlap with those of KBGS. Indeed, some patients diagnosed clinically with KBGS are found to have mutations in SETD5^40^. Motivated by this overlap, and by the effects of disease-causing TBLR1 mutations on ANKRD11/SETD5 binding, we investigated whether TBLR1 might coordinate the functions of these proteins. We find that TBLR1 forms a molecular “bridge” between ANKRD11 and SETD5, resulting in an assembly that, together with the NCoR complex, bears a striking resemblance to the yeast Set3 complex (SET3C) – a transcriptional regulator^41–45^. The relevance of this complex to neurodevelopmental disorders is further supported by our finding that pathogenic missense mutations in both ANKRD11 and SETD5 disrupt their association with SET3C. Moreover, *Setd5*^W834C^, a mutation engineered to specifically disrupt TBLR1 binding, resulted in severe developmental defects in mice, similar to those observed in *Setd5*-*null* animals. In mouse embryonic stem cells, this same mutation, and a pathogenic mutation in *Ankrd11*, give rise to highly correlated changes in gene expression. Furthermore, depletion of SET3C complex components in a human cultured cell line^46^ resulted in highly correlated changes in gene expression, further implicating SET3C in transcriptional regulation. Intriguingly, following perturbation of human SET3C subunits in this system, the most significantly upregulated genes were amongst the most transcriptionally active. Overall, our results reveal TBLR1 as a keystone of the transcriptional regulator SET3C, whose dysfunction underlies a family of prevalent neurodevelopmental disorders.

## Results

### Pathogenic TBLR1 mutations disturb MeCP2, SETD5 and ANKRD11 binding

To identify interaction partners affected by pathogenic mutations in TBLR1, we used mass spectrometry to compare proteins that co-immunoprecipitate (co-IP) with wild-type and mutant forms of TBLR1. We used CRISPR to inactivate the endogenous TBLR1 gene in Flp-In T-REx 293 cells (Extended Data Fig. 1a), and then transiently expressed wild-type or mutated TBLR1-mCherry in these cells. We observed that the G70D mutation^6^, which affects the N-terminal domain of TBLR1 – made up of a LisH domain and a putative F-box motif^47^ – disturbs the interaction of TBLR1 with ANKRD11 (Fig. 1a and 1b). In contrast, the D370N^33^ mutation within the C-terminal WDR domain of TBLR1 reduced the interaction with SETD5 (Fig. 1a and 1b). We decided to focus on ANKRD11 and SETD5, both of which have been reported to interact with the NCoR complex^35–37^, since patients with mutations in either of these genes often present with KBG syndrome or similar neurodevelopmental symptoms^39,40^.

**Figure 1.**
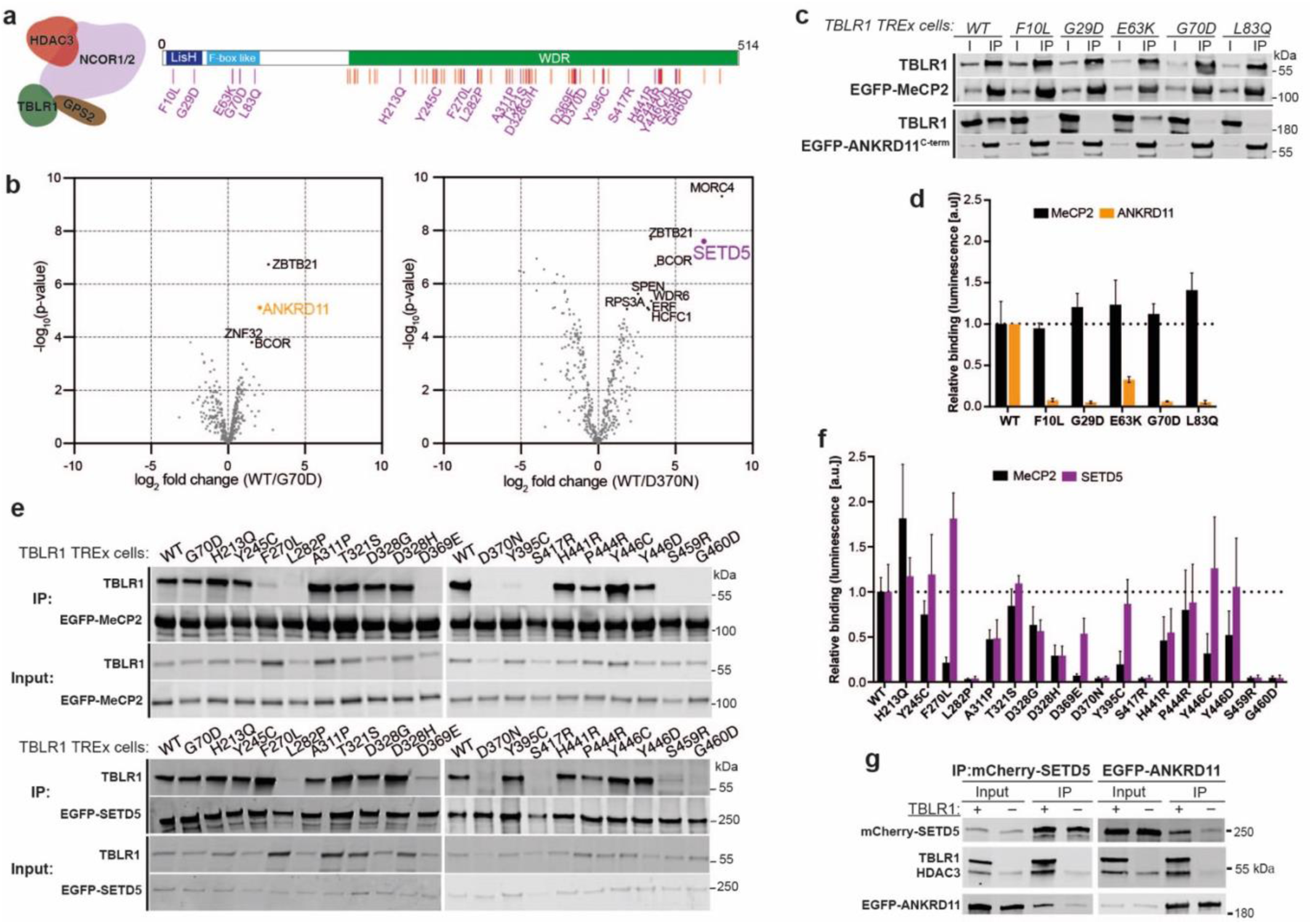
Pathogenic TBLR1 mutations disturb MeCP2, SETD5 and ANKRD11 binding. a, A schematic of the functional domains in TBLR1. The positions of pathogenic missense mutations from the ClinVar (likely pathogenic/pathogenic), DECIPHER (*de novo*, excluding benign mutations), and SFARI GENE (*de novo*) databases are indicated by vertical lines. The mutations we tested (purple) are labelled. **b,** A volcano plot showing enrichment of protein interactions detected by mass spectrometry after immunoprecipitation of wild-type (WT) or mutated (G70D or D370N) TBLR1-mCherry from TBLR1-null Flp-In™ T-REx™ 293 cells (n = 3 independent transfections per mutation)**. c,** Western blot for TBLR1 and EGFP following immunoprecipitation of EGFP-MeCP2 and EGFP-ANKRD11^C-term^ (ANKRD11 residues 1762-2663) from the indicated N-terminal TBLR1 mutant Flp-In™ T-REx™-293 cell lines. **d,** NanoLuc complementation assay using the indicated N-terminal mutant TBLR1-LgBiT constructs and either MeCP2-SmBiT or ANKRD11- SmBiT (n = 3-5 independent transfections per TBLR1 mutant). **e,** Western blot for TBLR1 and EGFP following immunoprecipitation of EGFP- MeCP2 or EGFP-SETD5 from the indicated WDR domain TBLR1 mutant Flp-In™ T-REx™ 293 cell lines. **f,** NanoLuc complementation assay using the indicated WDR domain mutant TBLR1-LgBiT constructs and either MeCP2-SmBiT or SETD5-SmBiT (n = 6-7 independent transfections per TBLR1 mutant). **g,** Western blot for TBLR1, HDAC3, mCherry and EGFP following immunoprecipitation of mCherry-SETD5 or EGFP-ANKRD11^C-term^ in either TBLR1-null or TBLR1^WT^ Flp-In™ T-REx™ 293 cells.

To test whether reduced binding to ANKRD11, SETD5 and/or MeCP2 is a general feature of pathogenic TBLR1 mutations, we created 24 TBLR1 mutant cell lines by integrating TBLR1 cDNAs at the specific FRT site-containing locus in our *TBLR1*-null Flp-In T-REx 293 line (Extended Data Fig. 1a and 1b). We then transiently expressed GFP-tagged MeCP2, SETD5 or ANKRD11 and assayed their interaction with TBLR1 mutants by IP with an antibody against GFP followed by western blotting with an antibody against TBLR1. Focussing first on pathogenic missense mutations in the N-terminal domain of TBLR1 (Fig. 1a), all five changes drastically reduced binding to ANKRD11 (Fig. 1c). In contrast, binding to MeCP2, which is known to interact with the C-terminal WDR domain of TBLR1, was unaffected by N-terminal mutations in TBLR1 (Fig. 1c). To validate the effects of TBLR1 mutations on binding to ANKRD11, we used a NanoLuc luciferase protein fragment complementation assay^48,49^. In agreement with the co-IP data, N-terminal TBLR1 mutations strongly reduced binding to ANKRD11, without affecting interaction with MeCP2 (Fig. 1d). The N-terminus of TBLR1 mediates self-association, and also incorporation into the NCoR complex^47^. To test whether pathogenic missense mutations in this region of TBLR1 might affect these interactions, FLAG-TBLR1 and either wild-type or mutant mCherry-tagged TBLR1 were transiently expressed together in TBLR1-null cells. IP of wild-type or mutant TBLR1-mCherry, followed by western blotting with FLAG antibodies revealed that TBLR1 self-association was not affected by these mutations (Extended Data Fig. 1c). Moreover, N-terminal mutations in TBLR1 did not markedly reduce TBLR1 association with the NCoR complex, as shown by Western blotting with antibodies against NCoR complex components NCoR1 and HDAC3 (Extended Data Fig. 1c). We conclude that pathogenic mutations in the N-terminus of TBLR1 interfere specifically with its interaction with ANKRD11.

We next assayed the effect of pathogenic missense mutations in the WDR domain of TBLR1 on binding to SETD5 and MeCP2. In co-IP experiments, approximately half of the eighteen WDR domain mutations tested strongly reduced binding to both SETD5 and MeCP2 (Fig. 1e), with the effects of mutations on SETD5 binding being well-correlated with their effects on MeCP2 binding. WDR domain mutations, which strikingly reduced binding to SETD5 and/or MeCP2 as assessed by co-IP assays, also showed strong reductions in binding in the NanoLuc complementation assay (Fig. 1f). Furthermore, for all but two of the remaining TBLR1 WDR mutations (H213Q and T321S), the more quantitative NanoLuc assay revealed moderate reductions in binding that were not apparent in the co-IP experiments (Fig. 1f).

As altered stability can contribute to the dysfunction of proteins carrying disease-causing missense mutations^31,50^, we tested the effects of pathogenic TBLR1 mutations on TBLR1 protein levels (Extended Data Fig. 1d and 1e). All N-terminal mutant forms of TBLR1 were expressed at approximately the same level as the wild-type protein. In most cases, WDR domain mutations did not substantially affect TBLR1 expression levels either. However, we observed up to an approximately two-fold reduction in TBLR1 expression level for WDR domain mutations with the most severe effects on MeCP2 and SETD5 binding (Extended Data Fig. 1e). Of note, patients heterozygous for deletion mutations that completely remove TBLR1^4,5^, and therefore have a 50% dose of the protein, display relatively mild symptoms^33,51,52^, making it unlikely that the more severe pathology associated with some TBLR1 missense mutations is primarily driven by their effects on TBLR1 abundance. Instead, the results suggest that for the majority of disease-causing TBLR1 mutations, disruption of their interaction with MeCP2, ANKRD11 and/or SETD5 contributes to the associated pathology.

The effects of TBLR1 mutations on binding to ANKRD11 and SETD5 raised the possibility that TBLR1 serves as a molecular bridge joining these two proteins, whilst also functioning as an adaptor protein to connect ANKRD11/SETD5 with the rest of the NCoR complex. To test this, we performed co-IP assays of mCherry-SETD5 and GFP-ANKRD11 transiently expressed in *TBLR1*-null cells or control cells expressing wild-type TBLR1. In the absence of TBLR1, the ability of ANKRD11 and SETD5 to interact with each other, and with the HDAC3 subunit of the NCoR complex, was strongly reduced (Fig. 1g). Residual binding observed in these experiments may be due to low levels of TBL1X in our TBLR1-null cells. We conclude that TBLR1 is necessary for robust interaction between ANKRD11, SETD5 and the rest of the NCoR complex.

### The complex between ANKRD11, SETD5 and NCoR resembles the yeast SET3 complex

The data presented point to the existence of a complex between SETD5, ANKRD11 and the other subunits of NCoR. It has previously been noted that SETD5 resembles the SET domain protein SET3 that is part of the yeast histone deacetylase-containing SET3 complex (SET3C)^37,41^. Further parallels exist between the subunits of SET3C and the components of the mammalian complex under study. Amino acid sequence conservation is limited, but most protein domains within subunits of the yeast complex have matching counterparts in mammals (Extended Data Fig. 2). These include an ankyrin-like repeat domain in HOS4 (similar to ANKRD11), a catalytically inactive SET domain in SET3 (similar to SETD5), a LisH domain and a WDR domain in SIF2 (similar to TBLR1), a SANT domain in SNT1 (similar to NCOR1) and a histone deacetylase domain in HOS2 (similar to HDAC3). Based on extensive domain conservation, we consider that the ANKRD11/SETD5/NCoR complex represents a mammalian ortholog of SET3C.

### Disease-causing missense mutations in ANKRD11 disturb SET3C assembly

Our data demonstrate that many pathogenic mutations in TBLR1 disturb ANKRD11/SETD5 binding, and so disruption of SET3C may represent an underlying cause of the associated disorders. According to this hypothesis, mutations in SETD5 and ANKRD11 that affect SET3C assembly would be expected to cause disease. Considering first ANKRD11, the C-terminal domain contains a cluster of pathogenic KBGS mutations (Fig. 2a), and is reported to act as transcriptional repressor in reporter assays^53^. IP of transiently expressed GFP-tagged ANKRD11 truncations followed by western blot analysis revealed that a 292 amino acid C-terminal fragment of ANKRD11 is sufficient for binding to the NCoR complex. In contrast, other regions of ANKRD11 did not show detectable interaction with NCoR (Fig. 2b). Five pathogenic C-terminal mutant forms of ANKRD11 identified in patients with developmental delay and cognitive impairments^54^ (S2475P, R2512Q, E2522K, R2523W and L2605R) were transiently expressed in HEK293 cells as GFP-fusions, and IPs were assayed by mass spectrometry. The mutants all showed a substantial reduction in binding to both NCoR complex subunits and to SETD5 (Fig. 2c). Reduced NCoR binding by mutant ANKRD11 was confirmed by IP of GFP-ANKRD11 and western blotting with antibodies against NCoR complex components (Fig. 2d). Thus, a putative globular domain at the C-terminus of ANKRD11 binds to the TBLR1-containing NCoR complex (Fig. 2e), and mutations disrupting this interaction lead to KBGS. This finding is consistent with the idea that assembly into SET3C is essential for ANKRD11 function.

**Figure 2.**
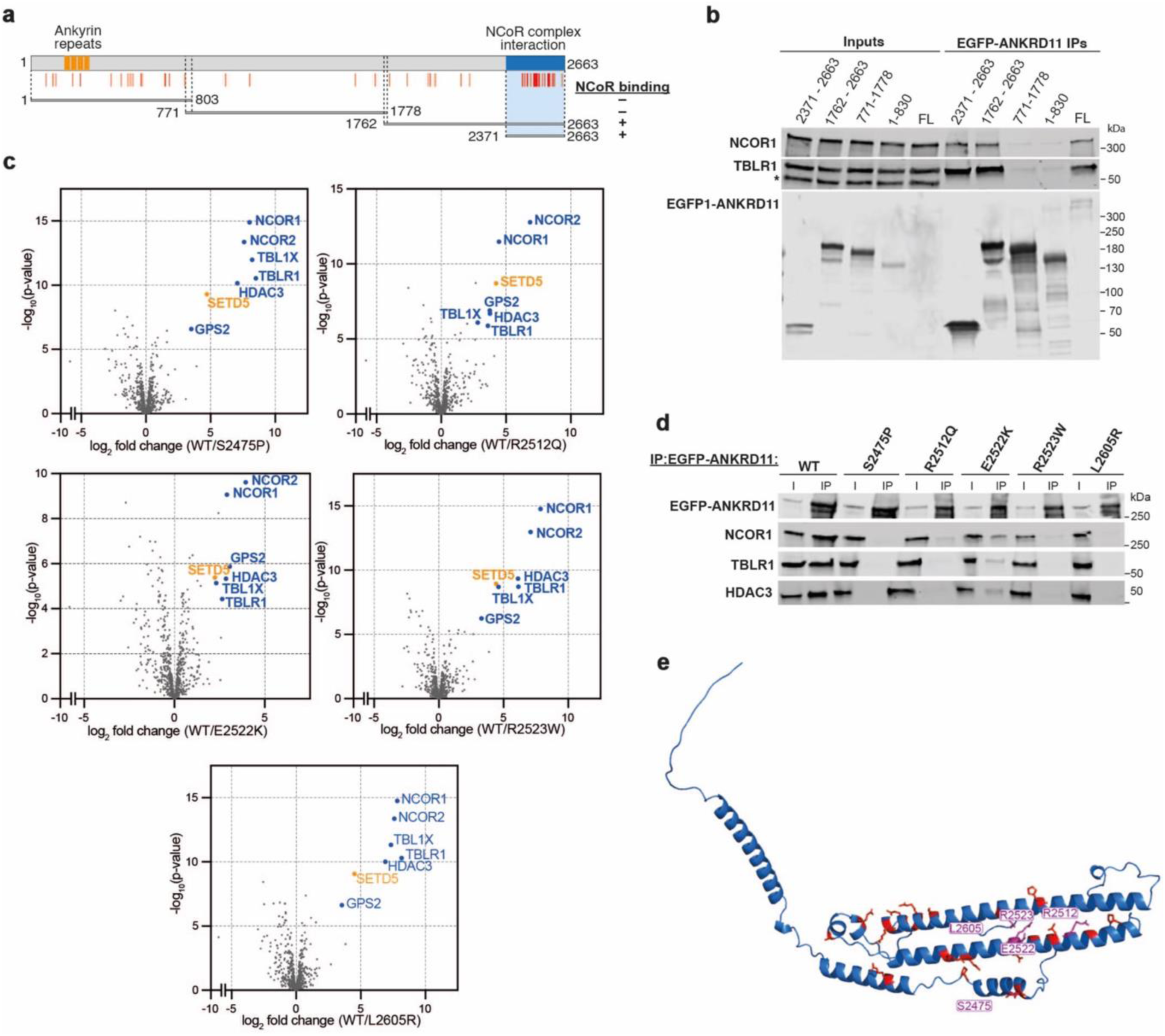
Disease-causing missense mutations in ANKRD11 disturb SET3C assembly. a, A schematic of the functional domains in ANKRD11. The positions of pathogenic missense mutations from the ClinVar (likely pathogenic/pathogenic), DECIPHER (*de novo*, excluding benign mutations), and SFARI GENE (*de novo*) databases are indicated by vertical lines. Truncation mutations tested are annotated. **b,** Western blot for TBLR1, NCoR1 and EGFP following immunoprecipitation of EGFP- ANKRD11 full length (FL) or truncation constructs from TBLR1^WT^ Flp-In™ T-REx™ 293 cells. **c,** Volcano plots showing enrichment of protein interactions detected by mass spectrometry following immunoprecipitation of wild-type (WT) EGFP-ANKRD11 or missense mutant constructs from HEK293T cells (n = 3 independent transfections per mutant). NCoR complex core components are indicated in blue. SETD5 is in orange. **d,** Western blot for EGFP, NCoR1, TBLR1 and HDAC3 following immunoprecipitation of wild-type (WT) or indicated missense mutant EGFP- ANKRD11 expressed in TBLR1^WT^ Flp-In™ T-REx™ 293 cells. **e,** AlphaFold 3 predicted structure of a C-terminal globular domain in ANKRD11. Positions of pathogenic mutations are highlighted in red with those used in this study indicated in purple.

### An MeCP2-like motif in SETD5 mediates its incorporation into SET3C

To locate the region of SETD5 responsible for incorporation into SET3C, we tested a series of mCherry-tagged truncations that progressively remove C-terminal regions of SETD5 (Fig. 3a). Co-IP assays showed that 1-846 still bound to the NCoR complex, whereas the 1-486 and 1-223 truncations did not bind (Fig. 3a and Extended Data Fig. 3a). Examination of the SETD5 amino acid sequence between 486 and 846 revealed the motif PLKKWKS, which closely resembles the core TBLR1-binding NID of MeCP2 (PIKKRKT)^20,21^ (Fig. 3b). This offered a potential explanation for the similar effect of different TBLR1 mutations on binding to SETD5 and MeCP2 (Fig. 1e and 1f). To test whether the SETD5 NID-like motif can interact with TBLR1, we transiently expressed wild-type or mutated GFP-SETD5 in Flp-In T-REx cells expressing wild-type TBLR1. The W834C mutation was chosen, since a mutation at the corresponding position in MeCP2 (R306C) disrupts binding to TBLR1 and causes Rett syndrome (Fig. 3b)^20,21^. After IP of GFP-SETD5, mass spectrometry showed that the W834C mutation severely disrupts the association of SETD5 with components of SET3C, including ANKRD11 (Fig. 3c). The disruptive effect of the SETD5 W834C mutation on NCoR complex and ANKRD11 binding was confirmed by co-expressing mCherry-SETD5 with GFP- ANKRD11^C-term^, followed by anti-mCherry IP and western blotting with antibodies against TBLR1, HDAC3 and GFP (Fig. 3d). Pull-down assays using a biotin-tagged SETD5 NID peptide, and extracts from either wild-type mouse brain or cells transiently expressing GFP-ANKRD11^C-term^, further demonstrated that this region of SETD5 is sufficient for SET3C incorporation (Fig. 3e and Extended Data Fig. 3b).

**Figure 3.**
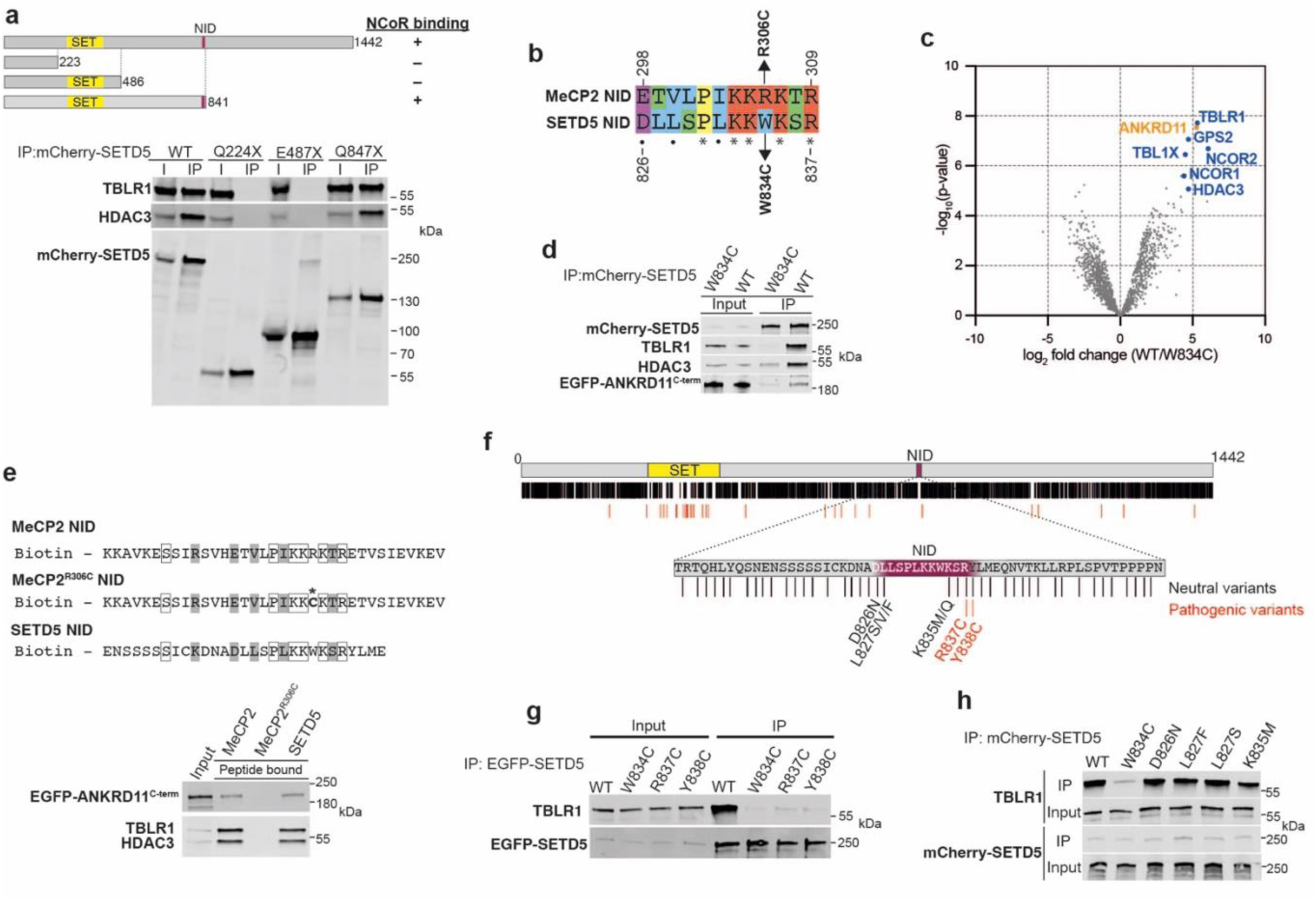
An MeCP2-like motif in SETD5 mediates its incorporation into SET3C, mutation of which causes disease. a, Western blot for TBLR1, HDAC3 and mCherry following immunoprecipitation of wild-type (WT) mCherry-SETD5 or indicated truncation mutants from TBLR1^WT^ Flp-In™ T-REx™ 293 cells. The location of the putative SETD5 NID (NCoR interaction domain) is indicated on the schematic below. **b,** Sequence alignment of MeCP2 and SETD5 NIDs (NCoR interaction domains). Amino acids are coloured based on their properties (Clustal X). asterisks = same amino acid, circles = similar amino acid. The positions of the R306C RTT-causing mutation in MeCP2 and the equivalent W834C mutation in SETD5 are indicated. **c,** Volcano plots showing enrichment of protein interactions detected by mass spectrometry following immunoprecipitation of wild-type (WT) or W834C mutant EGFP-SETD5 expressed in TBLR1^WT^ Flp-In™ T-REx™ 293 cells (n = 3 independent transfections). NCoR complex core components are labelled in blue. ANKRD11 is labelled in orange. **d,** Western blot for TBLR1, HDAC3, mCherry and EGFP following immunoprecipitation of wild-type (WT) or W834C mutant mCherry-SETD5 after co- transfection of TBLR1^WT^ Flp-In™ T-REx™ cells with mCherry-SETD5 and EGFP-ANKRD11^C-term^ (residues 1762-2663). **e,** Western blot for EGFP, TBLR1 and HDAC3 after peptide pull-downs from EGFP-ANKRD11^C-term^ transfected TBLR1^WT^ Flp-In™ T-REx™-293 cell lysates. Sequences of the biotinylated peptides are shown with identical (white box) and similar (grey) amino acids indicated. * = R306C Rett Syndrome causing mutation in MeCP2 which is known to abolish TBLR1 binding. **f,** A schematic of the functional domains in SETD5. The positions of neutral missense mutations from gnomAD 4.1 (excluding variants also in ClinVar) are indicated by black vertical lines. The positions of pathogenic missense mutations from the ClinVar (likely pathogenic/pathogenic), DECIPHER (*de novo*, excluding benign mutations), and SFARI GENE (*de novo*) databases are indicated by red vertical lines. **g,** Western blot for TBLR1 and EGFP following immunoprecipitation of wild-type (WT) or pathogenic mutant (R837C and Y838C) EGFP-SETD5 expressed in TBLR1^WT^ Flp-In™ T-REx™ 293 cells. The W834C mutation is used as a negative control. **h,** Western blot for TBLR1 and mCherry following immunoprecipitation of wild-type (WT) and mutant mCherry-SETD5 expressed inTBLR1^WT^ Flp-In™ T-REx™ 293 cells. The mutations tested are from gnomAD v2.1.1, v3.1.2 and ExAC (excluding mutations also in ClinVar).The W834C mutation is used as a negative control.

### Disease-causing missense mutations in SETD5 disturb SET3C assembly

Two pathogenic missense mutations in SETD5, R837C and Y838C, are located close to the NID (Fig. 3f). To test the effect of these variants on TBLR1 binding, we performed IP of wild-type or mutant GFP-SETD5 followed by western blotting with antibodies against TBLR1. Both mutations drastically reduced the interaction of SETD5 with TBLR1 (Fig. 3g), suggesting that loss of this interaction might contribute to the molecular pathology in these patients. Consistent with the NID being a key functional motif in SETD5, this region of the protein is relatively depleted of neutral variants in individuals with no evidence of developmental delay (gnomAD). Neutral changes of this kind are present relatively close to the NID at residues D826, L827 and K835 (Fig. 3f), which, if the NID is critical for SETD5 function, should not affect SETD5 binding to the NCoR complex. IP of four gnomAD mutants expressed as GFP-SETD5 fusions, followed by western blotting with antibodies against TBLR1 confirmed that none had any detectable effect on binding (Fig. 3h). Together, these observations offer strong support for the idea that SETD5 function relies on the ability of its NID to interact with TBLR1, and that genetic disease associated with SETD5 mutations can involve loss of this interaction.

### SET3C is required for embryonic development

To determine whether normal mouse development also requires incorporation of SETD5 into the SET3C complex, we generated mice with a W834C mutation in *Setd5* (Extended Data Fig. 4a). Heterozygous *Setd5^W834C/+^* animals were viable and fertile but had somewhat reduced body weight – including kidney weight (Extended Data Fig. 4b) – and length compared to wild-type littermates (Fig. 4a). Brain weight was similar to controls for *Setd5^W834C/+^* mice resulting in an increased brain to body weight ratio (Fig. 4a). These features closely match phenotypes previously described in mice heterozygous for a *Setd5*-null allele^35,55^. Homozygosity for this null mutation leads to embryonic lethality^37^. Similarly, when heterozygous *Setd5^W834C/+^* animals were inter-crossed, only 3.6% of pups were homozygotes (n = 4/110), which is well below the expected Mendelian ratio of 25% (Fig. 4b). Therefore, failure of mutant W834C SETD5 to join the SET3C complex resembles the effects of a null mutation in the mouse *Setd5* gene, including, in the great majority of cases, pre-natal lethality.

**Figure 4.**
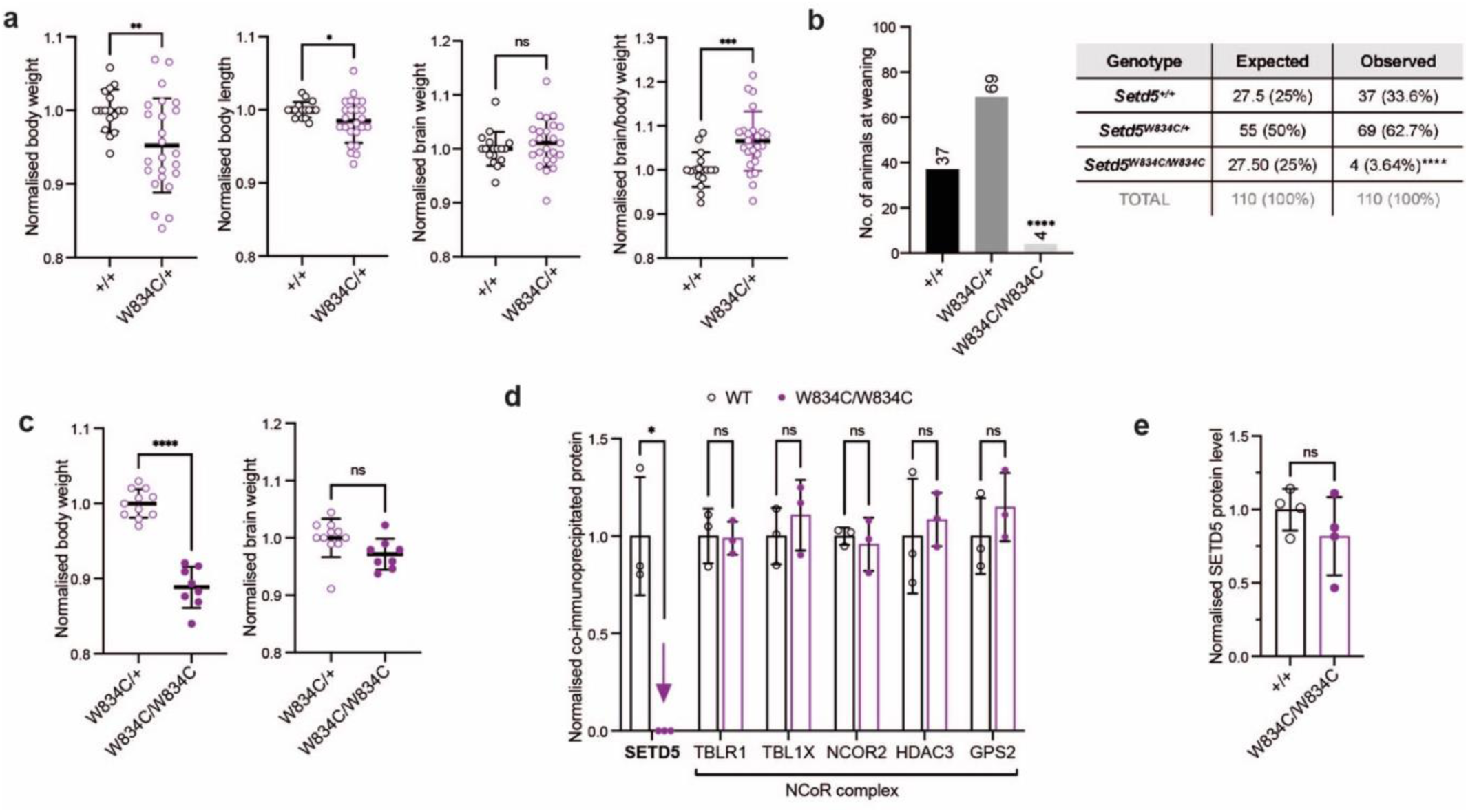
SET3C is required for embryonic development. a, Body and brain weight, body lengths and brain/body weight ratio of heterozygous *Setd5^W834C/+^* mice (n = 24) and their wild-type *Setd5^+/+^* littermates (n = 17). Values were normalised to the average of the wild-type littermates of the same sex. Means and standard deviations are plotted. Statistical significance was calculated using Welch’s unpaired t-test (body weight **p = 0.0028, body length *p = 0.0291, brain weight ^ns^p = 0.3475, brain/body weight ***p = 0.0004). **b,** Numbers of *Setd5^W834C/W834C^*, *Setd5^W834C/+^* and *Setd5^+/+^* offspring at weaning from *Setd5^W834C/+^*X *Setd5^W834C/+^* crosses. Statistical significance was calculated using a Chi squared test (p < 0.0001). **c,** Body and brain weights of homozygous *Setd5^W834C/W834C^*mice (n = 9) and heterozygous *Setd5^W834C/+^* (n = 6) littermates. Values were normalised to the average of the *Setd5^W834C/+^* littermates of the same sex. Means and standard deviations are plotted. Statistical significance was calculated using an unpaired t-test (body weight ****p < 0.0001, brain weight ^ns^p = 0.1593). **d,** Mass spectrometry quantification of NCoR complex components and SETD5 bound following immunoprecipitation with an antibody against NCoR1 from extracts from wild-type (WT) vs. W834C/W834C homozygous mouse brains. Statistical significance was calculated using unpaired t-tests (SETD5 *p = 0.004660). **e,** Mass spectrometry quantification of SETD5 protein levels in wild-type (+/+) vs. W834C/W834C homozygous mouse brain extracts (n = 3 brains per genotype). Statistical significance was calculated using unpaired t-test (^ns^p = 0.276069).

The surviving homozygous *Setd5^W834C/W834C^* mice had body weights that were more severely reduced relative to controls, whereas brain weight once again remained similar to littermates (Fig. 4c). Using brain nuclear extracts from these mice, we confirmed that the W834C knock-in mutation in endogenous *Setd5* prevents binding to NCOR1 without affecting SETD5 protein expression levels (Fig. 4d and 4e). As homozygotes were fertile, we were able to ask whether they were rare survivors of a developmental bottleneck or if they might have been rescued by a secondary genetic change. Compatible with the latter hypothesis, a cross between homozygous *Setd5^W834C/W834C^* and heterozygous *Setd5^W834C/+^* mice produced a substantially higher proportion of homozygous pups (Extended Data Fig. 4c). Testing this possibility and identifying the cause of suppression will require further work. Taken together, the mouse data demonstrate that a single amino acid change preventing SETD5 from assembling into SET3C has severe phenotypic consequences. We conclude that assembly of SET3C is a critical aspect of the molecular function of SETD5.

### The integrity of SET3C is required for transcriptional regulation

The resemblance of mammalian SET3C to yeast SET3C suggests that its role may also be to modulate gene expression. To test this possibility, we performed RNA-seq on mouse embryonic stem cells (ESCs) homozygous for ANKRD11 or SETD5 missense mutations that disrupt their association with SET3C. A by-product of generating homozygous *Setd5* W834C ESCs was a clone homozygous for an in-frame deletion of amino acids K833 to K835 of SETD5. This mutation severely disrupted the ability of SETD5 to bind to TBLR1 and only moderately affected SETD5 protein levels (Extended Data Fig. 5a and 5b). We therefore analysed both W834C and ΔKWK mutant ESCs by RNA-seq (two independent W834C clones and one ΔKWK clone). *Setd5* mutant ESCs showed 2844 differentially expressed genes (DEGs) compared to the wild-type controls with similar numbers of genes up- and down-regulated (1588 up, 1256 down) (Fig. 5a). In the case of ANKRD11, we generated three independent clones of ESCs with a missense mutation equivalent to the KBG syndrome mutation E2522K. This mutation is present in the Yoda mouse line^56^ where homozygosity is associated with *in utero* lethality, and heterozygosity causes behavioural and neurological abnormalities^57^. *Ankrd11^yod/yod^* ESCs had a greater number of DEGs, again with similar numbers of up and down regulated genes (11063 total DEGs; 5774 up and 5289 down) (Fig. 5a). The Yoda mutation, unlike the W834C mutation or ΔKWK in SETD5, substantially destabilises the ANKRD11 protein in both ESCs and in mouse brain (Extended Data Fig. 5c). This, or the presence of potentially redundant orthologous proteins, may explain its greater effect on transcription (see Discussion). Importantly, the transcriptional changes associated with these mutations are highly correlated (2057 total shared DEGs; 1078 up and 880 down) (Fig. 5b and c), supporting the view that ANKRD11 and SETD5 operate in the same pathway. Shared DEGs include many involved in organogenesis, including differentiation of muscle, heart, bone and kidney (see top 25 GO terms in Extended Data Fig. 5d). These altered transcriptomes suggest that the mutant cells have a propensity to differentiate.

**Figure 5.**
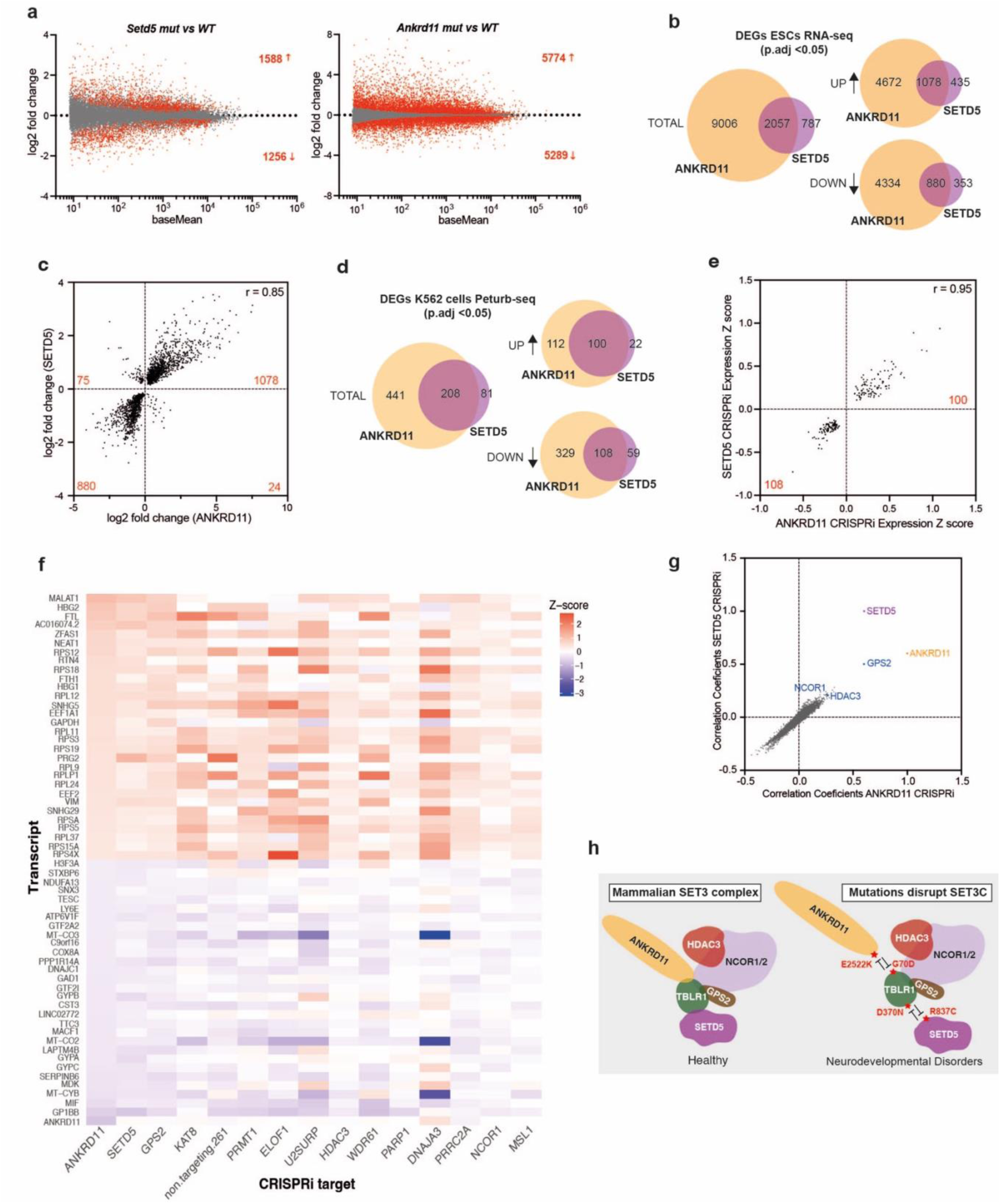
The integrity of SET3C is required for transcriptional regulation. a, MA plots of gene expression changes in *Setd5* and *Ankrd11* mutant mouse embryonic stem cells. Genes with significantly increased or decreased expression levels (p adj < 0.05) are shown in red (n = 3 independent clones for each genotype). **b,** Venn diagrams showing the overlap of differentially expressed genes (DEGs) in *Setd5* and *Ankrd11* mutant ESCs. Separate diagrams are shown for total DEGs, up-regulated DEGs and down-regulated DEGs. **c**, Scatterplot showing gene expression changes in *Ankrd11* and *Setd5* mutant mouse embryonic stem cells (R = 0.85). Points correspond to genes which are differentially expressed in both mutants (p adj < 0.05). **d,** Venn diagrams showing the overlap of differentially expressed genes (DEGs) following CRISPRi against SETD5 or ANKRD11 in K562 cells. Separate diagrams are shown for total DEGs, up-regulated DEGs and down-regulated DEGs. **e,** Scatterplot showing differentially expressed genes in K562 cells following CRISPR interference against ANKRD11 or SETD5. **f,** Heatmap showing gene expression changes after CRISPRi against ANKRD11 and the twelve other proteins with the highest correlation coefficients with ANKRD11. The top 30 up- and down-regulated genes in ANKRD11 CRISPRi (ordered by Z-scores) are shown. **g,** Scatterplot showing correlation coefficients of genome-scale CRISPRi scRNA-seq with CRISPRi against ANKRD11 and SETD5. **h,** Schematic of the proposed model of the mammalian SET3 complex. Patient mutation in SET3C components disrupt the assembly of the complex and cause neurodevelopmental disorders.

Since many DEGs shared by the mutant ES cell lines were related to differentiation, it seemed likely that these reflect destabilization of the stem cell state rather than primary effects of SET3C- deficiency on gene expression. In an attempt to reduce the downstream consequences of altered cell states and to test the effect of knocking down other SET3C subunits, we drew upon a Perturb-seq dataset derived from the human erythroid cell line K562 in which single-cell RNA-seq was used to assay the effects of genome scale CRISPR interference-mediated perturbations^46^. The gene expression changes caused by depletion of ANKRD11 or SETD5 were once again highly correlated (Fig. 5d and 5e). Indeed, they were better correlated with each other, than with any of the other approximately 11,000 perturbations tested (Fig. 5f and 5g). Furthermore, depletions of the NCoR subunits NCoR1, GPS2 or HDAC3 also produced transcriptional changes that were well-correlated with those observed when ANKRD11 or SETD5 are knocked-down (Fig. 5f and 5g). This data strongly suggests that ANKRD11, SETD5 and other components of SET3C function together to affect transcription. The most significantly dysregulated genes in Perturb-seq included ribosomal protein genes, haemoglobin, ferritin, vimentin, eosinophil major basic protein, and a set of abundantly expressed long non-coding RNAs (Fig. 5f), all of which were up-regulated. A common feature of these genes is that their mRNA transcripts are already abundant in unperturbed erythroid K562 cells. An intriguing possibility that requires future research is that SET3C preferentially restrains expression of highly transcribed genes.

## Discussion

Data from the 100k genome project places *ANKRD11*, *SETD5* and *MECP2* within the top ten most frequently mutated genes contributing to monogenic neurodevelopmental abnormalities^38^. Through analysis of patient mutations in transfection experiments, mutant cells in culture and mouse models, we pinpoint disruption of ANKRD11-TBLR1-SETD5 interactions as a common aetiology underlying these disorders (Fig. 5h). Previously reported SETD5 binding proteins include the NCoR complex^35,37^, the PAF complex^37^, ANKRD11^35^, G9a/GLP^58^ and BRD2^59^, and previously reported ANKRD11 interactions include p53^60^, HDAC3^36^, the p160 coactivator^36^, AIB1^61^, ADA3^62^, cohesin^63^ and the minor spliceosomal protein 65K/RNPC3^64^ as well as self-association^65^. However, despite this information, it remained uncertain whether disruption of mechanisms involving these interactions leads to the pathology seen in SETD5 syndrome or in KBGS. Our data now suggest that brain function relies on the integrity of SET3C – the mammalian equivalent of the yeast SET3C complex. Strikingly, disruptions of different SET3C subunits can have similar effects on gene expression profiles. Thus, failure of SET3C-dependent transcriptional regulation is a shared feature of a set of distinct but related neurodevelopmental disorders.

Our findings help to explain why TBLR1 mutations are not associated with a single discrete clinical disorder. We find that TBLR1 has at least two clinically relevant functions (binding to MeCP2 and SET3C), and different mutations in TBLR1 disrupt these to different extents. For example, mutations in the TBLR1 N-terminal domain that prevent its association with ANKRD11 will disrupt the assembly of SET3C while preserving the ability of TBLR1 to serve as an adaptor for NCoR recruitment by MeCP2. Other mutations in the TBLR1 WDR domain interfere with both MeCP2 and SETD5 binding, and these might blend both RTT-like pathology and the pathology associated with SET3C dysfunction. There are additional reasons why mutations in TBLR1 might only rarely resemble RTT. TBLR1 is autosomal whereas MeCP2 is an X-linked gene and therefore subject to X chromosome inactivation. Classical RTT is the result of a mosaic brain with complete loss of MeCP2-dependent functions in approximately half of cells, the other half being functionally wildtype, whereas TBLR1 mutations would be associated with a molecular pathology that affects every cell. The relationship between different TBLR1 mutations and variable clinical outcomes might be further complicated by effects of TBLR1 mutations on its other binding partners. One candidate is ANKRD12, a paralog of ANKRD11 for which loss-of-function mutations are associated with poor verbal-numerical reasoning^66^. Also, MLL5, a paralog of SETD5, may be functionally redundant with SETD5. Yet another binding partner is USP44 – a ubiquitin protease that binds to the TBLR1 WDR domain^67^ and has been implicated in cognitive disability^68^. Finally, the implications of our observation that MeCP2 and SETD5 interact with the same surface of TBLR1 via a closely related amino acid motif are also uncertain. An intriguing possibility is that the two proteins compete for access to NCoR, suggesting that mutations in one may enhance or disrupt the interaction with the other, with potential phenotypic consequences.

Our findings indicate that SET3C may limit the transcription of highly expressed genes. This is consistent with a recent report that in *Drosophila* embryos, HDAC3 depletion upregulated genes which were already highly expressed in control tissues^69^. Work in yeast suggests that the SET3 complex transcriptionally suppresses sporulation genes^41^ and also spurious transcriptional initiation from cryptic promoters^42,70^. The yeast SET3 complex is reportedly targeted to the dimethylated form of lysine 4 of histone H3 via the PHD finger of its Set3 subunit^43^. However, although SETD5 is related to Set3, it does not contain a comparable PHD finger (unlike its mammalian paralog MLL5). *Drosophila* UpSET – an ortholog of SETD5 that also has a PHD finger – has been implicated in preventing the ectopic spread of active chromatin away from genes and into neighbouring flanking regions^71^. ChIP-seq from different groups suggests that SETD5 binds preferentially to transcription start sites^35^, or to active gene bodies^55^. Understanding SET3C function will require reconciliation of these observations via elucidation of the molecular determinants of SET3C binding to chromatin. Several interactions and domains could plausibly play a role in specifying genomic binding, including the PAF complex, the ankyrin repeat domain of ANKRD11, the SET domain of SETD5 (which contains several pathological missense mutations), and other subunits of SET3C. Recruitment via the ANKRD11 ankyrin repeats is an attractive hypothesis since they share significant homology with the ankyrin repeats of BARD1 – a domain that binds to the N-terminal tail of histone H4^72^. At present, however, the mechanism by which SET3C is recruited to chromatin remains uncertain.

The finding that mutations affecting these three genes affect a shared regulatory pathway involved in related neurological disorders is relevant to the quest for therapies. Most immediately, lessons learned about the treatment of patients with one SET3C-related disorder might help to to guide the management of patients with other related conditions. Looking further ahead, efforts to develop gene therapy as a treatment strategy for SETD5 syndrome or KBGS may be complicated by the large size of SETD5 and ANKRD11, due to the limited capacity of available viral vectors. One possible solution to this problem is genome editing, which is relatively indifferent to gene size – a technology that is rapidly developing. Another is to develop truncated versions of SETD5 and ANKRD11, which nevertheless retain all their critical domains and motifs, and so can function normally. The design of such constructs will require a complete inventory of the functional elements in SETD5 and ANKRD11. Just as reporting the key MeCP2-TBLR1 interaction^20,21^ paved the way for developing the miniaturised version of MeCP2^32^ that underlies the first RTT gene therapy clinical trial^73^, our findings on SET3C-related disorders may also hold translational potential.

## Methods

### Plasmids

The coding sequences of ANKRD11, SETD5, TBLR1 and/or MeCP2 were cloned into pEGFP/mCherry-C1, pmCherry-N1, p3xFLAG-CMV10 and/or pcDNA5/FRT/TO (Thermo, V652020). The NanoLuc LgBiT or SmBiT fragments were added to the C-terminus of coding sequences (see Table S1 for plasmid sequences). Truncation and missense mutation variants of these plasmids were produced by PCR and Gibson assembly or by site-directed mutagenesis (see Table S1 for primer sequences).

### Western blotting

Samples for western blots were either immunoprecipitation samples, peptide pull-down samples, input samples or whole-cell protein extracts. For whole-cell protein extracts: cell pellets were lysed in NE10 buffer (20 mM HEPES pH 7.9, 10 mM KCL, 1 mM MgCl2, 0.1% Triton X-100, 20% glycerol, 0.5 mM DTT, Benzonase 1000U/ml and protease inhibitors cocktail) and incubated at room temperature for 10 min. 2x SDS-PAGE sample buffer was added and solution vortexed, boiled for 3 min, snap frozen on dry ice then boiled for a further 3 min. Antibodies used for western blotting were as follows: ANKRD11 (Santa Cruz Biotechnology, sc-514916), HDAC3 (Abnova, H00008841- M02), HDAC3 (Abcam, ab32369), mCherry (Abcam, ab167453), NCoR1 (Cell Signalling Technology, 5948S), SETD5 (Santa Cruz, Sc-515645), GFP (Takara, 632592), GFP (Takara, 632381), SIN3A (Abcam, ab3479), TBLR1 (Santa Cruz Biotechnology, sc-100908), γ-tubulin (Sigma, T5326) and Histone H3 (Abcam ab1791). Western blots were imaged on the LI-COR® Odyssey CLx system and quantified using the Image Studio Lite software (LI-COR biosciences).

### Cell culture

Mouse embryonic stem cells (ESCs) (129/Ola E14TG2a) were cultured on gelatinized culture dishes and flasks in Glasgow MEM medium (GMEM) (Gibco 21710025) supplemented with recombinant mouse Leukemia Inhibitory Factor (LIF) (ESGRO), 10% FBS (Gibco), 1% Non-essential amino acids (Gibco)), 1% Sodium pyruvate (Gibco), and 0.1% β-mercaptoethanol (Gibco). HEK293 cells (ATCC, CRL-1573) were grown in DMEM (Gibco, 41966029) supplemented with 10% FBS (Gibco) and transfected with lipofectamine 2000 (Thermo). Flp-In™ T-REx™ 293 cells (Thermo, R78007) were cultured according to manufacturer’s protocol. Cells were harvested 24 hours after transfection and/or induction of protein expression and then stored at -80°C for future analysis.

*Setd5^W834C/+^* heterozygous ESCs were generated using two CRISPR-Cas9 guide RNAs (Table S1) targeting either side of exon 17 and a long single stranded DNA (lssDNA) template with the W834C mutation and silent PAM-abolishing mutations (Table S1). The ssDNA template was excised from a plasmid using the nicking endonucleases Nt.BspQI and Nb.BsmI, and purified using the LssODN Preparation Kit (Biodynamics Laboratory Inc., Tokyo, Japan). *Setd5^W834C/W834C^* homozygous ESCs were generated using a single CRISPR-Cas9 guide RNA (Table S1) and a 200 bp DNA oligonucleotide (IDT) containing the W834C mutation (Table S1). Targeted clones were identified and verified by XcmI restriction digest (the W834C mutation destroyed a XcmI site) and by sequencing.

*ANKRD11^E2523K/E2523K^* homozygous ESCs were made using a single CRISPR-Cas9 guide RNA (Table S1) and a 200 bp DNA oligonucleotide (IDT) containing the E2523K mutation and silent mutations within the seed region of the gRNA target DNA sequence (Table S1). Targeted clones were identified and verified by MseI restriction digest (inserted mutations introduce an MseI site) and by sequencing.

TBLR1 was knocked-out in Flp-In™ T-REx™ 293 cells using a CRISPR-Cas9 guide RNA (see Table S1) targeting a region just downstream of the ATG initiation codon. Subsequent sequencing identified clones which were compound heterozygous for frame shift mutations in this region.

Tetracycline-inducible expression constructs, derived from pcDNA5/FRT/TO (see Table S1), were inserted into the FRT site of Flp-In™ T-REx™ 293 cells according to the manufacturer’s instructions. Single-cell clones were verified by sequencing and single integration of the plasmid into the FRT site was confirmed by PCR and Southern blotting (see Table S1 for primers).

### Animals

Mice were bred in a specific-pathogen-free facility in individually ventilated cages, with wood chippings, tissue bedding and environmental enrichment and were given *ad libitum* access to food and water. Rooms were maintained on a 12-hour light/dark cycle at 45-65% humidity at 20-24°C. Procedures were carried out by certified persons, licensed by the UK Home Office and according to the Animals (Scientific Procedures) Act 1986.

The *Setd5^W834C^* mice were generated by injection of heterozygous *Setd5^W834C/+^* ESCs (see above) into E3.5 blastocysts obtained from C57BL/6J females after natural matings. Blastocysts were transferred to pseudo-pregnant recipient females and chimeric offspring were mated to C57BL/6J mice to establish the line. The *Ankrd11^Yod^* mice (C3H.Cg-Ankrd11^Yod^/H) were rederived from mouse sperm purchased from the EMMA mouse repository (EM:00380). Mouse genotyping was initially performed by PCR (See Table S1 for primers) and XcmI digest for *Setd5^W834C^* mice then by TransnetYX. *Ankrd11^Yod^* and Setd5^W834C^ mice were genotyped by TransnetYX (TransnetYX assays available upon request). Weight and length measurements and tissues for molecular analysis were taken from 6 week-old mice.

### NanoLuc assays

NanoLuc complementation assays were performed as described before^49^ except that full-length TBLR1 was used rather than the isolated WDR domain.

### Mass spectrometry

Samples were prepared for mass spectrometry either by in gel digestion as described elsewhere^74^ or (for the analysis of pEGFP-SETD5 IPs) by filter aided sample preparation as described elsewhere^75^. Immunoprecipitations were done in triplicate from 3 separate cell transfections per construct or brains from 3 separate animals per genotype and all samples were analysed by mass-spectrometry.

LC-MS analyses were performed on Orbitrap Fusion™ Lumos™ Tribrid™ Mass Spectrometer (Thermo Fisher Scientific, UK) using the Data Dependent Acquisition (DDA) mode and on Orbitrap Exploris 480™ on a Data Independent Acquisition (DIA) mode, both coupled on-line, to an Ultimate 3000 HPLC (Dionex, Thermo Fisher Scientific, UK). Peptides were separated on a 50 cm (2 µm particle size) EASY-Spray column (Thermo Scientific, UK), which was assembled on an EASY-Spray source (Thermo Scientific, UK) and operated constantly at 50°C. Mobile phase A consisted of 0.1% formic acid in LC-MS grade water and mobile phase B consisted of 80% acetonitrile and 0.1% formic acid. Peptides were loaded onto the column at a flow rate of 0.3 μl min^-1^ and eluted at a flow rate of 0.25 μl min^-1^ according to the following gradient: 2 to 40% mobile phase B in 150 min and then to 95% in 11 min. Mobile phase B was retained at 95% for 5 min and returned back to 2% a minute after until the end of the run (190 min).

For DDA, survey scans were recorded at 120,000 resolution (scan range 350-1500 m/z) with an ion target of 4.0e5, and injection time of 50ms. MS2 was performed in the ion trap at a rapid scan mode, with ion target of 2.0e4 and HCD fragmentation^76^ with normalized collision energy of 27. The isolation window in the quadrupole was 1.4 Thomson. Only ions with charge between 2 and 7 were selected for MS2. Dynamic exclusion was set at 60 s.

For DIA, MS1 scans were recorded at 120,000 resolution (scan range 350-1650 m/z) with an ion target of 5.0e6, and injection time of 20 ms. MS2 was performed in the orbitrap at 30,000 resolution with a scan range of 200-2000 m/z, maximum injection time of 55 ms and AGC target of 3.0E6 ions. We used HCD fragmentation with stepped collision energy of 25.5, 27 and 30. We used variable isolation windows throughout the scan range ranging from 10.5 to 50.5 m/z. Narrow isolation windows (10.5-18.5 m/z) were applied from 400-800 m/z and then gradually increased to 50.5 m/z until the end of the scan range. The default charge state was set to 3. Data for both survey and MS/MS scans were acquired in profile mode.

MaxQuant^77^ version 1.6.1.0 was used to process the raw files and search was conducted against *the Mus musculus* protein database (released in July 2017), using the Andromeda search engine^78^. For the first search, peptide tolerance was set to 20 ppm while for the main search peptide tolerance was set to 4.5 pm. Isotope mass tolerance was 2 ppm and maximum charge to 7. Digestion mode was set to specific with trypsin allowing maximum of two missed cleavages. Carbamidomethylation of cysteine was set as fixed modification and oxidation of methionine was set as variable modifications. Label-free quantitation analysis was performed using the MaxLFQ algorithm as described^79^. Absolute protein quantification was performed as described^80^. Peptide and protein identifications were filtered to 1% FDR.

The DIA-NN software platform^81^ version 1.8.1 was used to process the DIA raw files and search was conducted against the *Mus musculus* complete/reference proteome (Uniprot, released in July, 2017). Precursor ion generation was based on the chosen protein database (automatically generated spectral library) with deep-learning based spectra, retention time and IMs prediction. Digestion mode was set to specific with trypsin allowing maximum of two missed cleavages. Carbamidomethylation of cysteine was set as fixed modification. Oxidation of methionine, and acetylation of the N-terminus were set as variable modifications. The parameters for peptide length range, precursor charge range, precursor m/z range and fragment ion m/z range as well as other software parameters were used with their default values. The precursor FDR was set to 1%.

The mass spectrometry proteomics data have been deposited to the ProteomeXchange Consortium via the PRIDE partner repository with the dataset identifier PXD063846.

### Binding assays

Brains were subject to Dounce homogenization in ice-cold NE10 buffer (20 mM HEPES NaOH pH 7.5, 10 mM NaCl, 1 mM MgCl2, 0.1% Triton-X-100, 10 mM 2-mercaptoethanol and protease inhibitors) followed by 5 min centrifugation at 500 g. Insoluble material was resuspended in NE10 and treated for 5 min at 25°C with 250 units of benzonase (Merck, E1014). 5 M NaCl was then added to give a final NaCl concentration of 150 mM. After 20 min mixing, and 30 min 16,000 g centrifugation, all at 4°C, the supernatant was taken as starting material for immunoprecipitation or peptide pull-down assays. Cell extract were prepared by Dounce homogenization in NE10 before benzonase treatment (as above), adjusting the NaCl concentration to 150 mM, and 30 min centrifugation at 16,000 g at 4°C. Supernatants were then taken as starting material for immunoprecipitations.

For immunoprecipitation (IP) of GFP-SETD5 for mass spectrometry anti-GFP (Merck, 11814460001) was used. For IP of NCOR1 from mouse brain anti-NCOR1 (Cell Signalling Technology, 5948S) was used. GFP trap (Chromotek, GTA-20) and RFP trap (Chromotek, RTA-20) were used for all other IPs. IPs with GFP/RFP trap were performed as described previously^20^. IPs with anti-GFP and anti-NCOR1 were performed similarly except that 5 µg of antibody was first bound to protein G Dynabeads (Thermo, 10003D) by cross-linking by mixing for 30 min at 20°C in borate buffer (40 mM boric acid, 40 mM sodium tetraborate decahydrate) with 20 mM dimethyl pimelimidate. Bound complexes were eluted by boiling with SDS-PAGE sample buffer, except for the IP of GFP-SETD5 for mass spectrometry, which was eluted by mixing for 15 min at 60°C in 50 µl 0.1% Rapigest in 50 mM Tris-HCl pH 8.0. For peptide pull-down assays 50 µg of peptide with an N-terminal biotin tag (see Table S1 for peptide sequences) was bound to 10 µl streptavidin sepharose (Merck, GE17-5113-01). As with the IP assays, complexes were then precipitated from extracts, washed, and eluted with SDS-PAGE sample buffer.

### RNA-seq

Total RNA was isolated from *Setd5^W834C/W834C^*, *Setd5^+/+^, Ankrd11^yod/yod^* and *Ankrd11^+/+^* ESC clones (n = 3 separate clones per genotype and n = 3 separate outgrowths per clone) using TRIzol (Thermo, 15596026) followed by purification with the Direct-zol RNA Miniprep Plus kit (Zymo Research, R2070) according to manufacturer’s protocol. Genomic DNA contamination was removed with the Invitrogen™ DNA-*free*™ DNA Removal Kit (AM1906) and the remaining DNA-free RNA was tested for purity using PCR for genomic loci. Total RNA was tested on the 2100 Bioanalyzer (Agilent Technologies) to ensure high quality and quantified using a Qubit (Invitrogen). Drosophila S2 cells (5% total cell count) were added to each sample as a spike in before proceeding with RNA-extraction. ERCC RNA Spike-in control mix (Ambion) was added to RNA samples before library preparation and sequencing. Strand-specific library preparation with rRNA-depletion was performed by Novogene followed by whole-transcriptome sequencing using HiSeq X (Novogene Europe, UK). With minor modifications, differential gene expression analysis was performed as described previously^82^.

## Acknowledgements

We thank Jan Ure and Joe Mee (Centre for Regenerative Medicine, University of Edinburgh) for the Ju09 129/Ola E14TG2a mouse ESC line. We thank the Bioresearch and Veterinary Services (BVS) Central Transgenic Core (CTC) (University of Edinburgh) for generating the *Setd5^W834C^* mouse model and rederiving the *Ankrd11^Yod^* mouse line. This work was funded by the Simons Initiative for the Developing Brain [SFARI-529085]. A Wellcome Investigator Award [107930] supports the work of A.P.B. This work was supported by funding for the Wellcome Discovery Research Platform for Hidden Cell Biology [226791] and we gratefully acknowledge support from the Proteomics core.

---

**Extended Data Figure 1:**
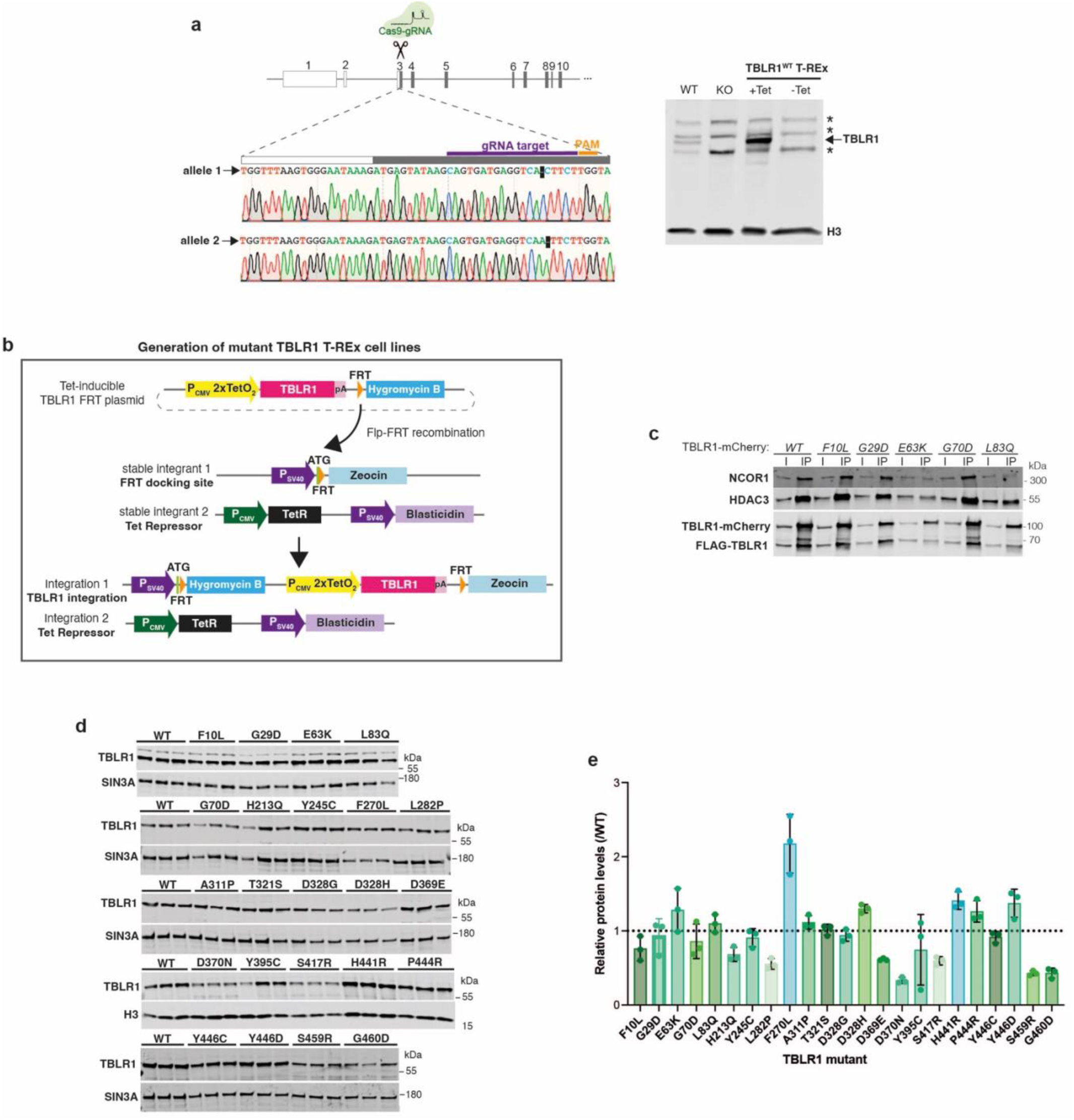
Pathogenic mutations in TBLR1 do not substantially affect protein stability. a, Left: Targeting strategy used to knock-out *TBL1XR1* in Flp-In™ T-REx™-293 cells and the sequences of the two targeted alleles with 1 bp deletions. Right: Western blot for TBLR1 and histone H3 using extracts from untargeted wild-type cells (WT), TBLR1 knock-out cells (KO) and KO cells with a TBLR1^WT^ cDNA under a tetracycline inducible promoter (TBLR1^WT^ T-Rex) inserted into the FRT site, treated with (+) or without (–) tetracycline. *Non-specific binding of TBLR1 antibody. H3 serves as a loading control. **b,** A schematic of Flp-recombinase mediated transgene insertion into the FRT site of Flp-In™ T-REx™-293 cells. **c,** Western blot for NCoR1, HDAC3, mCherry and FLAG following immunoprecipitation of wild-type (WT) or indicated mutated forms of TBLR1-mCherry from TBLR1^WT^ Flp-In™ T-REx™ 293 cells that were co-transfected with TBLR1-mCherry and FLAG-TBLR1. **d,** Western blot for TBLR1 and SIN3A/H3 using extracts from tetracycline-induced wild-type (WT) and indicated mutant TBLR1 Flp-In™ T-REx™ 293 cell lines (n = 3 independent tetracycline inductions for each mutation). SIN3A or Histone H3 serves as loading controls. **e,** Quantification of Western blots showing TBLR1 protein level relative to WT in the indicated mutant TBLR1 mutant Flp-In™ T-REx™-293 cell lines.

**Extended Data Figure 2:**
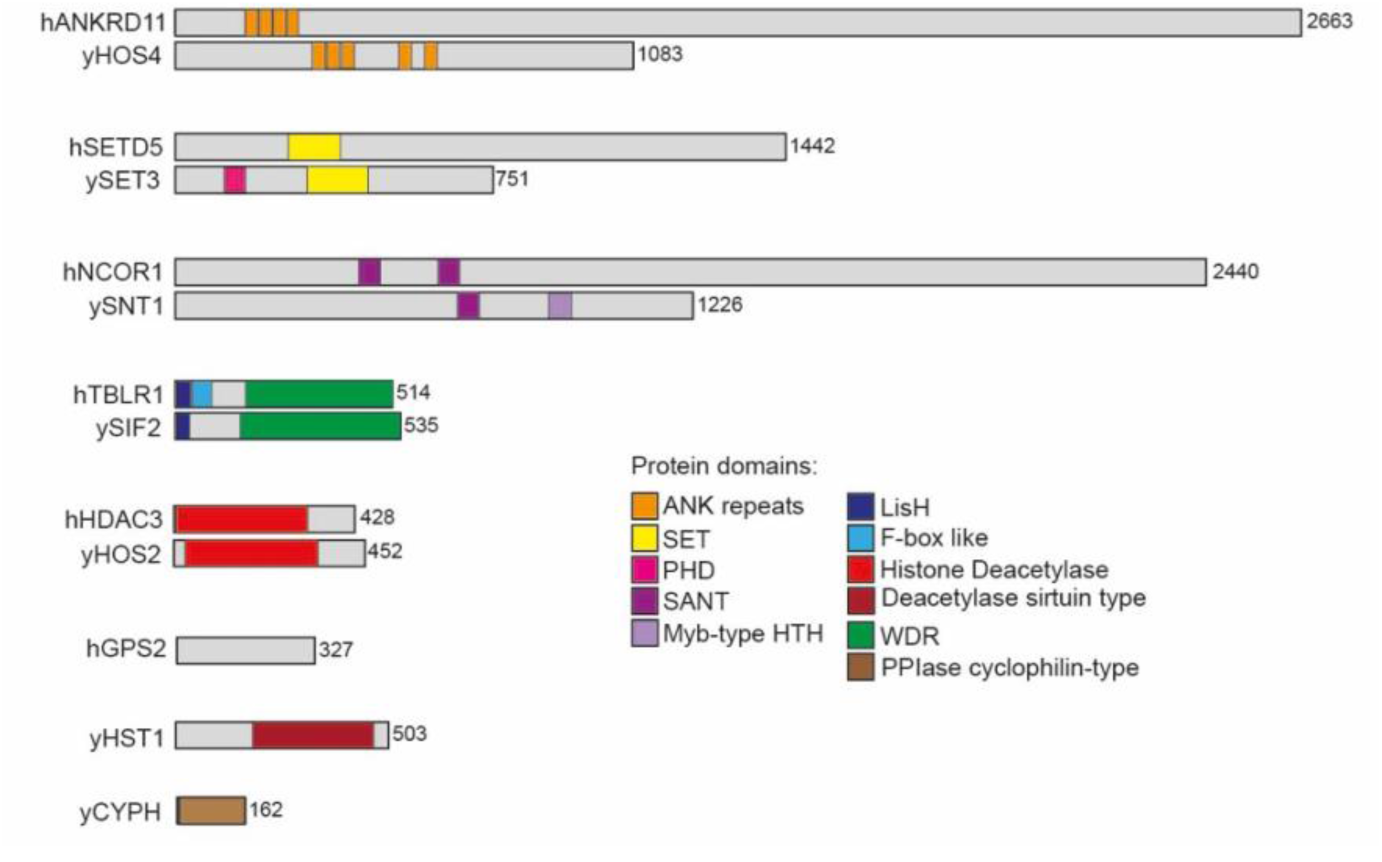
The complex between ANKRD11, SETD5 and NCoR resembles the yeast SET3 complex. A schematic showing domain structures of yeast (y) SET3 complex components and the analogous proteins in the human (h) assembly.

**Extended Data Figure 3:**
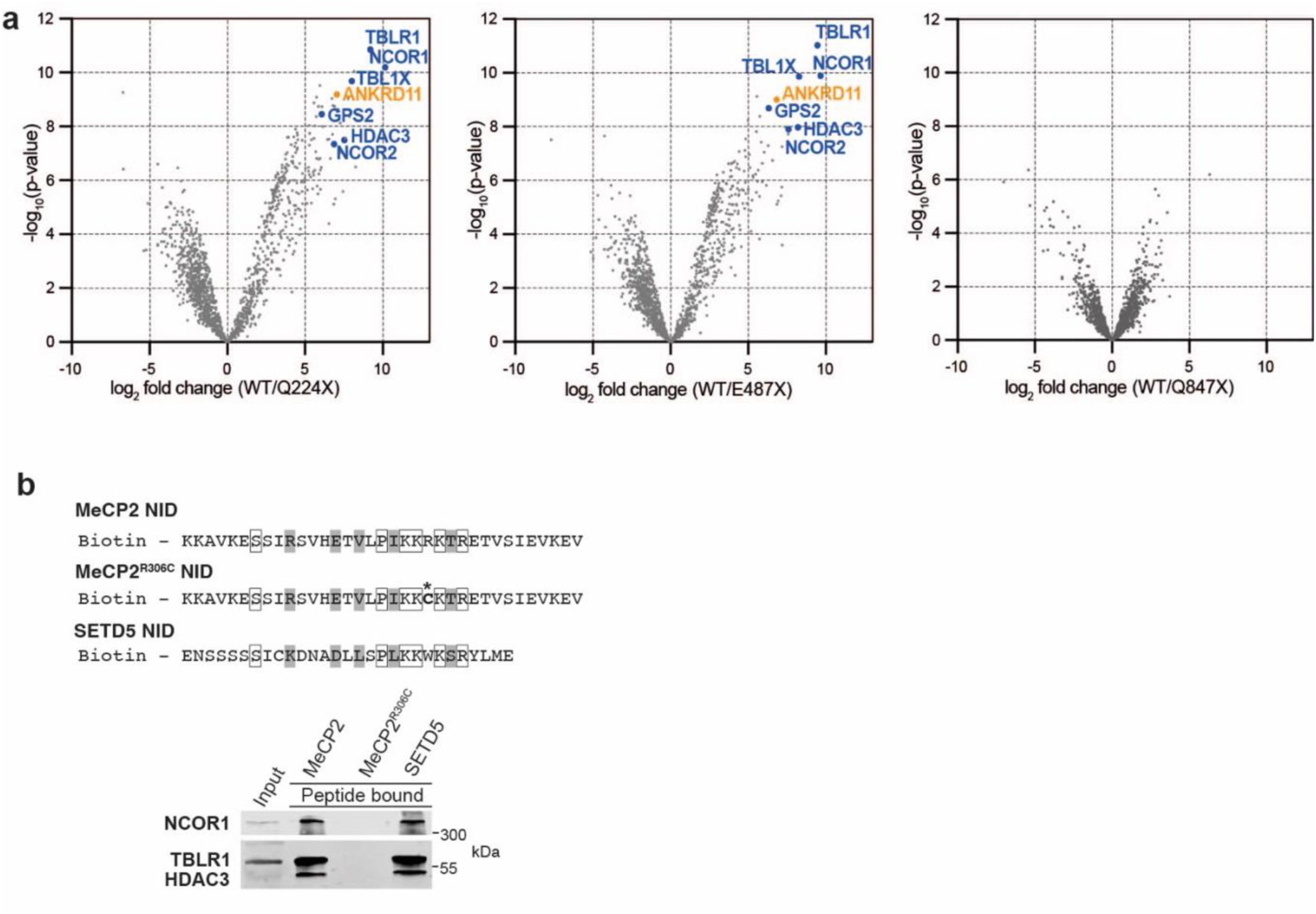
The NID motif in SETD5 mediates the interaction with SET3C. a, Volcano plots showing enrichment of protein interactions detected by mass spectrometry following immunoprecipitation of wild-type (WT) EGFP-SETD5 or indicated truncation constructs expressed in TBLR1^WT^ Flp-In™ T-REx™ 293 cells (n = 3 independent transfections). NCoR complex core components (blue) and ANKRD11 (orange) are labelled. **b,** Western blot for NCoR1, TBLR1 and HDAC3 after peptide pull- downs from wild-type mouse brain extracts. Sequences of the biotinylated peptides are shown with identical (white box) and similar (grey) amino acids indicated. Asterisks = R306C Rett Syndrome causing mutation in MeCP2 which is known to abolish TBLR1 binding.

**Extended Data Figure 4:**
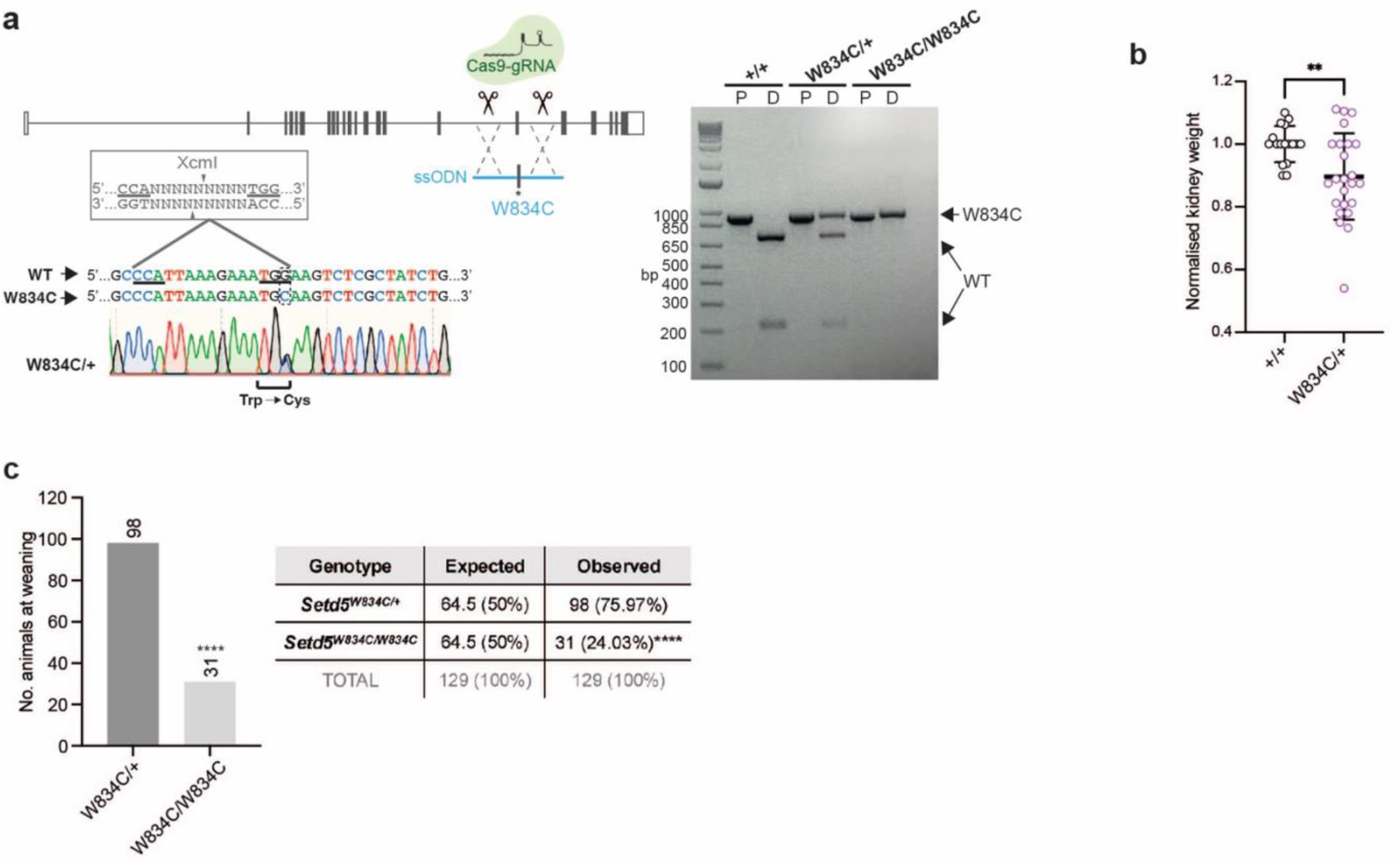
Generation and characterisation of the *Setd5^W834C^* mouse model. a, Left: Targeting strategy to generate the heterozygous *Setd5^W834C/+^* embryonic stem cells used to make mice. A Sanger sequencing chromatogram of a PCR amplicon of the targeted region from a heterozygous sample is shown. The W834C mutation destroys an XcmI restriction enzyme site. Right: The genotyping strategy used to distinguish *Setd5^+/+^, Setd5^W834C/+^* and *Setd5^W834C/W834C^* mice. Gel electrophoresis of PCR samples before and after XcmI digestion (P = PCR, D = digested). **b,** Kidney weight of heterozygous *Setd5^W834C/+^* mice (n=24) and their wild-type *Setd5^+/+^* littermates (n=17). Values were normalised to the average of the wild-type littermates of the same sex. Means and standard deviations are plotted. Statistical significance was calculated using Welch’s unpaired t-test (kidney weight **p = 0.0023). **c,** Numbers of *Setd5^W834C/W834C^* and *Setd5^W834C/+^* offspring of weaning age from *Setd5^W834C/W834C^* X *Setd5^W834C/+^* crosses. Statistical significance was calculated using a binomial test (****p < 0.0001).

**Extended Data Figure 5:**
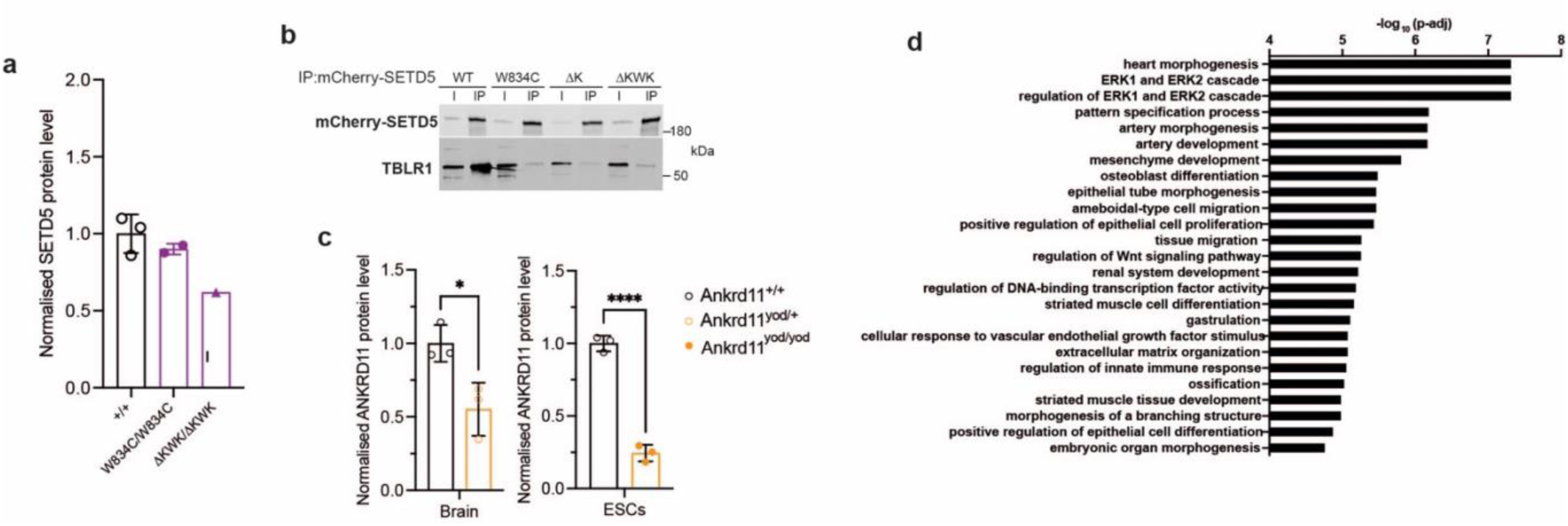
The SETD5 NID mutations destroy binding to TBLR1 without substantially affecting protein stability. a, Mass spectrometry analysis of SETD5 protein levels in extracts from wild-type, W834C homozygous and ΔKWK homozygous mouse embryonic stem cells. **b,** Western blot for TBLR1 and mCherry after immunoprecipitation of wild-type (WT) and mutant mCherry-SETD5 expressed in TBLR1^WT^ Flp-In™ T-REx™ 293 cells. **c,** Mass spectrometry quantification of ANKRD11 protein levels in nuclear extracts from Yoda heterozygous mouse brain and whole cell extracts from Yoda homozygous embryonic stem cells (ESCs). Statistical significance was calculated using unpaired t-test (Brains *p=0.0242, ESCs ****p <0.0001). **d,** GO term enrichment analysis of shared differentially expressed genes in *Setd5* and *Ankrd11* mutant mouse embryonic stem cells.

